# A simple device to immobilize protists for electrophysiology and microinjection

**DOI:** 10.1101/859413

**Authors:** Anirudh Kulkarni, Nicolas Escoubet, Léa-Laetitia Pontani, Alexis Michel Prevost, Romain Brette

## Abstract

We present a simple device to mechanically immobilize motile cells such as ciliates and flagellates. It can be used in particular for intracellular electrophysiology and microinjection. A transparent filter with holes smaller than the specimen is stretched over an outlet. A flow is induced by either a peristaltic pump or a depressurized tank, mechanically entraining cells to the bottom, where they immobilize against the filter. The cells swim again freely as soon as the flow is stopped. We demonstrate the device by recording action potentials in Paramecium and injecting a fluorescent dye in the cytosol.

## INTRODUCTION

*Paramecium* can swim at speeds exceeding ten times its body length per second. Thus, a key requirement to experimentally manipulate it is to immobilize it without damaging it. Classically, intracellular electrophysiology in ciliates such as *Paramecium* and *Tetrahymena* was performed with the hanging droplet method (Hennessey and Kuruvilla, 1999; Naitoh and Eckert, 1972). A specimen is picked with as little fluid as possible and placed hanging below a coverslip; the later use of inverted microscopes allowed the droplet to be placed on top of the coverslip (Houten, 1979; Valentine and Van Houten, 2016). When water evaporates, the cell is captured by surface tension. A hooked pipette is then gently but swiftly raised into the cell, effectively pinning it to the coverslip. The cell is then quickly covered by the bath before it dries out completely. This technique requires substantial dexterity. An additional difficulty is that this technique provides no electrical signal for impaling the cell, which must then entirely rely on visual inspection. A less common strategy is to catch the swimming organism with a suction pipette (Jonsson and Sand, 1987). For microinjection, the standard method consists in covering the specimen with oil, removing fluid with a needle until the cell is immobilized, then performing the microinjection and releasing the cell (Beisson et al., 2010).

Here we present a simple device to mechanically immobilize while providing an electrical signal. A transparent filter with holes smaller than the cells is placed at the bottom of the device, immersed in the bath. Fluid is then removed from the bottom using a peristaltic pump or a depressurized reservoir. In a few seconds, cells are immobilized against the filter. A pipette can then be inserted into the cell. If the pipette is filled with a conducting solution, successful impalement is indicated by a drop in measured potential. We demonstrate the use of the device by recording action potentials in *Paramecium Tetraurelia* using two electrodes, and microinjecting Alexa Fluor into the cytosol.

## MATERIALS AND METHODS

### *Paramecium* culture and manipulation

Cultures of *Paramecium tetraurelia* (obtained from Éric Meyer, Institut de Biologie, École Normale Supérieure, Paris, France) were maintained by reinjecting each week 1 mL of culture inoculated with *Klebsiella pneumoniae* into 5 mL of Wheat Grass Powder (WGP) buffer supplemented with 1 μL of beta-sitosterol. Cultures are kept at room temperature. Prior to each experiment, the culture is filtered through a LCH Pure SN30 non-woven sterile swab, and cells are washed and concentrated in a clean buffer (the extracellular solution used for electrophysiology, see below) using gravitaxis (Naitoh and Eckert, 1972). Indeed, Paramecia tend to accumulate at the top of any aqueous solution. Once a droplet of culture (typically 600 µL) is placed in a narrow neck volumetric flask, one can then recover a concentrated population at the top of the flask.

### Device fabrication and assembly

The device was engineered to provide immobilization of *Paramecia* by suction on the filter. It was fabricated with a combination of laser-cutting and micro-milling techniques. It consists of two thin Plexiglas plates (lower plate thickness ∼2.7 mm, upper plate thickness ∼1.3 mm) that sandwich a filter once they are tightly screwed together (Fig. 1A). For our experiments, we have used in particular transparent engineered Whatman Cyclopore polycarbonate membranes (diameter 25 mm, pore diameter 12 μm). Note that before assembly, the filter is first wet with water to ensure good adhesion with the device. The upper plate was laser-cut with a circular and centered hole (diameter 5 mm) in order to form a pool-like structure in which Paramecia can swim freely, once apposed against the lower plate. A mesa-like structure (diameter 4 mm, height 100 µm) was micro-milled in the center of the lower plate using a three-axis commercial desktop CNC Mini-Mill machine (Minitech Machinary Corp., USA) as shown in Figures 1B and 1C. The mesa’s purpose is to allow for the stretching of the filter just like a thin membrane is stretched on a drum. On the mesa structure, microfluidic channels (width 300 µm, depth 100 µm) were then micro-milled with the cross-like geometry shown in Figures 1a and 1b. Finally, a small through-hole (diameter 300 µm) was drilled in the center of the mesa, and eventually enlarged (diameter 600 µm) on 2 mm from the lower side of the plate. This allowed to insert a small metallic tube (tubing connector SC23/8, Phymep) that acts as a fluid outlet and to which external tubing can be easily connected and with which suction can be applied (Fig. 1C). The geometry of the microfluidic pattern was chosen to prevent any local bending of the filter while the mesa structure avoids larger height fluctuations, upon suction. Both plates were drilled with through-holes (diameter 2.2 mm) so screws (2 mm in diameter) combined with bolts could be used to assemble both parts of the device.

**Figure 1.**
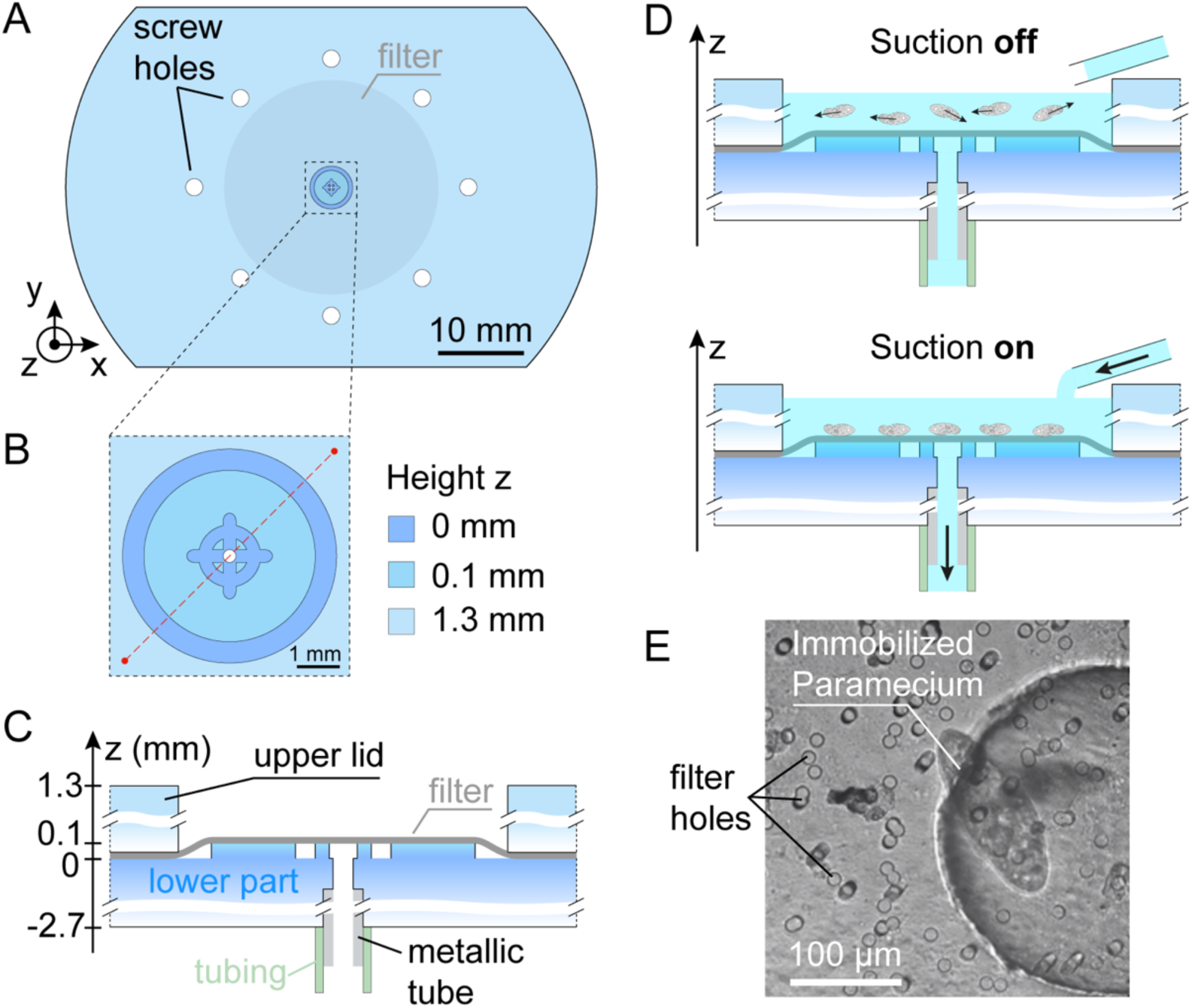
Sketch of the immobilization device. (A) Top view of the device. The filter (shaded grey disk) is sandwiched between an upper lid and a lower part all made in Plexiglas, and tightened together with 8 screws. (B) Close-up on the centered microfluidic mesa-like structure. The z=0 height origin is arbitrarily taken at the base of the mesa-like structure. (C) Lateral view along the red dashed-line cut in (B). (D) Principle of the immobilization process. Without suction (upper panel), *Paramecia* in the centered pool swim freely. Once suction is switched on (lower panel), *Paramecia* are immobilized against the filter. Their bathing liquid is pumped using either a peristaltic pump or a depression tank. The liquid is reinjected (when using the pump) or supplemented (when using the depression tank) in the pool to maintain the level of the bath constant. (E) Image in transmission of a single *Paramecium* immobilized with the current device (top view).

### Principle of the immobilization process

*Paramecia* are immobilized by simply sucking the liquid bath through the filter. As shown in the upper panel of Fig. 1D, without suction, *Paramecia* swim freely. But, as soon as suction is switched on (Fig. 1D, lower panel), the resulting hydrodynamic flux in the bath immobilizes the *Paramecia* against the filter.

### Pumping methods

The bathing liquid is pumped using either a peristaltic pump or a depression tank. It is reinjected in the pool to maintain a constant volume of the bath. In the case of a peristaltic pump, the tube of a Gilson Minipulse 3 pump is first filled with the medium (see extracellular solution in Electrophysiology, below). When using a depression tank, the device outlet is connected to a sealed glass jar with two entries, one for the tube from the device, and another one used to depressurize it to about −150 mbar. To apply a controlled pressure, we use a microfluidic flow controller (OB1 Mk3, Elveflow). However, a simple syringe can also be used to lower the pressure in the jar. Volume of the bath is maintained by being supplemented with a gravity perfusion system at a flow rate of 5 mL/min, while the excess solution is drained from the top by a peristaltic pump. In this way, it is possible to use the immobilization device together with the perfusion system, for example to exchange solutions while the *Paramecium* is immobilized, as shown on Supplementary Figure 1.

**Supplementary Figure 1.**
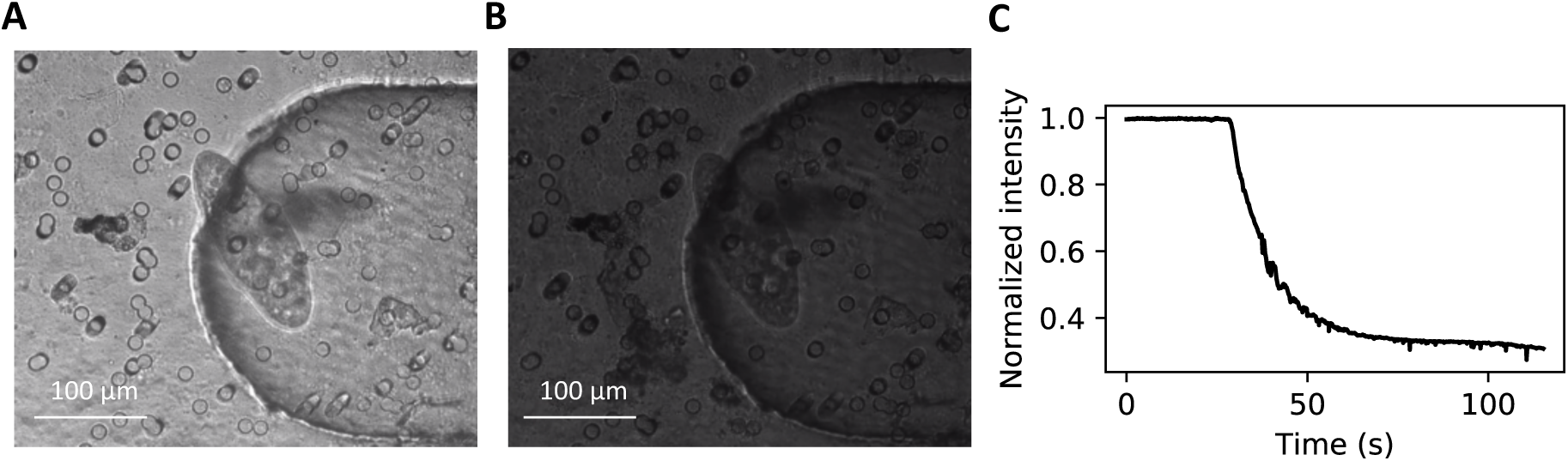
Solution exchange while a Paramecium is immobilized. (A) Paramecium immobilized against the filter by depression. (B) Immobilized Paramecium at the end of the solution exchange. (C) Change in normalized image intensity after the bath is replaced by a solution stained with Copper chlorophyllin, using a gravity perfusion system. Normalized intensity decreases at an initial rate of 0.04 / s, which is the expected value for an exchange flow rate of 5 mL/min and a 2 mL bath volume.

### Electrophysiology

For all experiments, we used a controlled extracellular solution consisting of 1 mM CaCl2, 4 mM KCl and 1 mM Tris-Hcl, pH=7.2. Microelectrodes of resistance ≈ 50 MΩ were pulled with a single step from standard wall borosilicate capillary glass with filament (OD 1 mm, ID 0.5 mm, Harvard Apparatus) using a micropipette puller (P-1000, Sutter Instrument). They were filled with a 1M KCl solution using a MicroFil non-metallic syringe needle (MF 34G-5, World Precision Instruments).

Custom Python programs (https://github.com/romainbrette/clampy) are used to control the analog-digital acquisition board (USB-6343, National Instruments) connected to the amplifier (Axoclamp 2B, Axon Instruments) operating at a sampling frequency of 40 kHz. After cell immobilization, the microelectrode is lowered into the cell until the measured potential drops by about 20 mV. The procedure is repeated with a second electrode. The pump or depression is then stopped. Square current pulses of amplitude 500 pA and duration 100 ms are then injected to tune the amplifier’s capacitance neutralization circuit.

### Microscopy

We image *Paramecium* using an upright microscope (LNScope, Luigs & Newmann) with two objectives, an air 20x objective (SLMPLN Plan Achromat, Olympus) used to locate cells, and a water immersion 40x objective (LUMPLFLN, Olympus) with DIC contrast enhancement for electrophysiology and microinjection. For visualization and recording, we use a high speed and high sensitivity CCD camera (Lumenera Infinity 3-6UR, 2752 × 2192 pixels^2^, 8 or 14 bit depth, 27 frames/s at full resolution). For fluorescence measurements, the setup is illuminated with a CoolLED pE-300 ultra combined with a Cy3 filter.

### Microinjection

Glass microinjection pipettes are pulled to an estimated outer diameter of 0.7-0.9 µm in one step using the same pipette puller as described above. The back of the pipette is connected to a microfluidic flow controller (OB1-Mk3, Elveflow) and controlled with ESI software (Elveflow Smart interface). The baseline pressure is set to 5 mbar, such that there is no net flow through the micropipette. *Paramecia* are injected with a solution containing 60 µM Alexa-594 fluorophore dye and 20 mM KCl, by applying a 100 ms long pulse at a pressure of 300 mbar.

## RESULTS AND DISCUSSION

### Immobilization

*Paramecia* swimming in a large drop are placed over the device. A downward flow can then be induced by two means. In the first configuration, a peristaltic pump draws fluid from the bath and pours it back at the top (see Movie 1, Supplemental Material). When the flow rate is greater than about 0.7 mL/min, *Paramecia* are pulled down and immobilized against the filter typically after a few seconds. Although *Paramecia* cannot swim, their cilia still beat. When the pump is stopped, *Paramecia* immediately swim away from the filter. Note that in practice, *Paramecia* can be immobilized by one or several holes constraining them in vertical or horizontal positions respectively. The hole diameter of the filter has thus to be smaller than the considered cell size.

In an earlier version of the device, the filter moved in the vertical direction by about 30 µm when the pump was turned on, as it pulled on it. To solve this issue, the filter is put on a slightly raised platform (see Methods), so that it gets stretched when the upper lid is screwed over the bottom part of the device. No measurable movement is then observed when the pump is turned on.

Since the peristaltic pump can introduce a periodic pulsation of Paramecia’s vertical positions, we also implemented a second configuration in which downward flow is induced by a negative pressure. In this configuration, the outlet is connected to a sealed reservoir. When the reservoir is depressurized at about −150 mbar, Paramecia are immobilized against the filter (see Movie 2, Supplemental Material). This pressure difference imposes a flow rate of about 0.7 mL/min into the reservoir, as in the first configuration. To maintain the liquid bath surface level in the pool, we use a gravity-based perfusion system that yields a flow rate of 5 mL/min, while the excess fluid is removed with a peristaltic pump. Perfusion can be used simultaneously with the depression; Movie 3 and Supplementary Figure 1 show a solution exchange while *Paramecium* is immobilized by depression.

### Electrophysiology

After immobilization, a pipette can be lowered into the cell. Figure 2A shows a *Paramecium* impaled with two sharp microelectrodes. Impalement is facilitated by the fact that, in contrast with the droplet technique, an electrical signal is available while the electrode is lowered. Indeed, entry of the microelectrode into the cytosol is witnessed by a voltage drop (Figure 2B and Movie 4). Once both microelectrodes are in place, the pumping flow is stopped. Figure 2C shows action potentials recorded by one electrode in response to current steps injected through the other electrode. If the pump is left running, the pulsation can be observed on the cell’s membrane but it usually does not impact the measured membrane potential (Movie 5). However, in one case we observed transient hyperpolarizations synchronized with the pulsation, indicative of mechanosensitive responses (Machemer and Deitmer, 1985). Therefore, it is advisable to switch off the pump or to avoid the pulsation by using the depression configuration.

**Figure 2.**
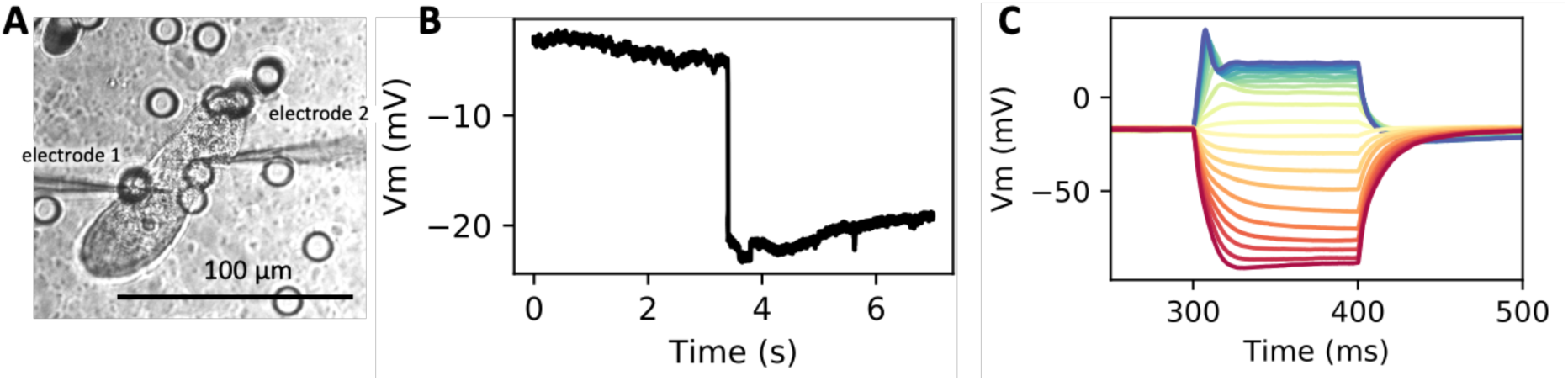
Electrophysiology on immobilized Paramecium. (A) Immobilized Paramecium impaled by two microelectrodes. (B)Membrane potential measured while the electrode is descended into the cell. Successful impalement is signaled by a sudden voltage drop. (C) Action potentials recorded in response to current steps of intensity −4 nA (red) to 4 nA (blue), by steps of 400 pA.

### Microinjection

Next, we perform a microinjection of a fluorescent solution in the cytosol of *Paramecium*. Since our specimens display green autofluorescence (Wyroba et al., 1981), we chose the red fluorophore Alexa Fluor-594 (Figure 3 and Movie 6, Supplemental Material). While the pump is still running, the fluorophore is injected by pressure (Figure 3A) and then removed. Figure 3B shows the fluorescent *Paramecium* a few minutes after microinjection. Noticeably, it swims normally once immobilization is stopped and retains its fluorescent content.

**Figure 3.**
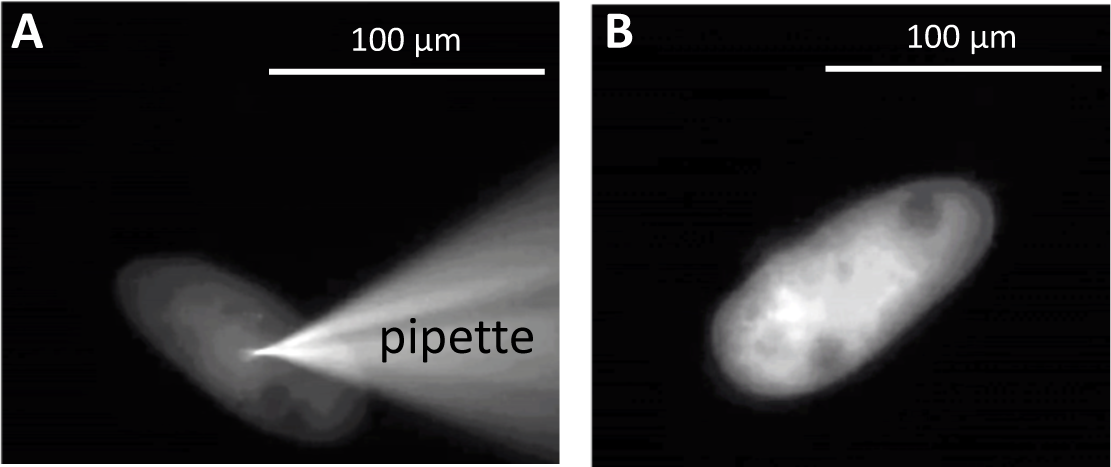
Microinjection of a fluorescent probe in an immobilized Paramecium. (A) Paramecium impaled by a microinjection pipette that contains Alexa Fluor-594. (B) Snapshot of a freely swimming Paramecium a few minutes after injection.

## Discussion

The device presented here was designed to ease the manipulation of motile *Paramecium* for both intracellular electrophysiology and microinjection measurements. Traditional methods mostly relied on trapping *Paramecium* in microdroplets. Two typical configurations were used, either trapping a single *Paramecium* in an aqueous droplet immersed in oil (Beisson et al., 2010), or confining it in an evaporating water droplet (Naitoh and Eckert, 1972). In the latter case, the time window during which one can approach a micropipette before *Paramecium* dies is very narrow, which leads to high failure rates. The former case is not adapted to electrophysiology because the micropipette tip gets contaminated with oil. In contrast, our method is easy to implement, highly reproducible, inexpensive and does not alter *Paramecium*’s viability. In particular, the immobilization can be obtained with any device that imposes a fluid flow such as peristaltic pumps, pressure controllers or syringe pumps. An additional benefit is that an electrical signal is available during the procedure, allowing one to verify the proper insertion of microelectrodes. Finally, our device allows for a straightforward medium exchange and is thus appropriate for easy drug testing on Paramecium.

One thus expects this device to be useful to efficiently trap any other type of motile protists or microorganisms provided that their typical size remains larger than the size of the filter holes. Beyond electrophysiology and microinjection, it may also allow imaging live cells over long periods of times, such as the sexual cycle. In the future, this immobilization technique could be straightforwardly automated by controlling the pump or using solenoid valves, which could allow complete automation of an electrophysiological or microinjection experiment.

## Supporting information

Reversible immobilization of Paramecia with a peristaltic pump.

Reversible immobilization of Paramecia with depression.

Solution exchange while a Paramecium is immobilized.

Impalement of immobilized Paramecium with two microelectrodes.

Intracellular voltage recording while the pump is switched on and off.

Microinjection of fluorescent Alexa in immobilized Paramecium.

Titles and captions of movies

## ACKNOWLEDGMENTS

The authors thank Eric Meyer for providing specimens and advice on culture and manipulation of *Paramecium*, and Martijn Sierksma for advice on electrophysiology.

## REFERENCES

Beisson J, Bétermier M, Bré M-H, Cohen J, Duharcourt S, Duret L, Kung C, Malinsky S, Meyer E, Preer JR, Sperling L. 2010. DNA Microinjection into the Macronucleus of Paramecium. Cold Spring Harb Protoc 2010:pdb.prot5364. doi:10.1101/pdb.prot5364

Hennessey TM, Kuruvilla HG. 1999. Chapter 17 Electrophysiology of Tetrahymena In: Asai DJ, Forney JD, editors. Methods in Cell Biology. Academic Press. pp. 363–377. doi:10.1016/S0091-679X(08)61543-5

Houten JV. 1979. Membrane potential changes during chemokinesis in Paramecium. Science 204:1100–1103. doi:10.1126/science.572085

Jonsson L, Sand O. 1987. Electrophysiological Recordings from Ciliates In: Scheving LE, Halberg F, Ehret CF, editors. Chronobiotechnology and Chronobiological Engineering, NATO ASI Series. Dordrecht: Springer Netherlands. pp. 432–434. doi:10.1007/978-94-009-3547-1_41

Machemer H, Deitmer JW. 1985. Mechanoreception in Ciliates In: Autrum H, Ottoson D, Perl ER, Schmidt RF, Shimazu H, Willis WD, editors. Progress in Sensory Physiology, Progress in Sensory Physiology. Berlin, Heidelberg: Springer Berlin Heidelberg. pp. 81–118. doi:10.1007/978-3-642-70408-6_2

Naitoh Y, Eckert R. 1972. Electrophysiology of Ciliate ProtozoaExp. in Physiol. and Biochem. pp. 17–31.

Valentine MS, Van Houten JL. 2016. Methods for Studying Ciliary-Mediated Chemoresponse in Paramecium. Methods Mol Biol Clifton NJ 1454:149–168. doi:10.1007/978-1-4939-3789-9_10

Wyroba E, Bottiroli G, P G. 1981. Autofluorescence of axenically cultivated Paramecium aurelia. Acta Protozool 20:165–170.

