## Supplementary material for "A simple device to immobilize protists for electrophysiology and microinjection": Titles and captions of movies

### Movie legends

***Movie 1.*** *Reversible immobilization of Paramecia with a peristaltic pump. The pump is switched on at t = 10 s and switched off at t = 40 s.*

***Movie 2.*** *Reversible immobilization of Paramecia with depression. Depression is on between t = 10 s and t = 40 s.*

***Movie 3.*** *Solution exchange while a Paramecium is immobilized. A stained solution (Copper chlorophyllin) replaces the clear extracellular solution.*

***Movie 4.*** *Impalement of immobilized Paramecium with two microelectrodes. The electrodes are filled with 1 M KCl and connected to an electrophysiological amplifier. A voltage drop appears when the electrode is inserted into the cell.*

***Movie 5.*** *Intracellular voltage recording while the pump is switched on and off. The pump is switched on at t = 20 s, then is switched on and off every 10 s. The pump pulsation is visible as movements of Paramecium, with no apparent change in membrane potential.*

***Movie 6.*** *Microinjection of fluorescent Alexa in immobilized Paramecium.*
